# Can an original be found? Mitochondrial species identity does not predict nuclear genome similarity in the photosymbiotic jellyfish *Cassiopea andromeda* and *C. xamachana*

**DOI:** 10.64898/2026.08.26.747389

**Authors:** Kaden Muffett, Megan Sporre, Marta Mammone, Maria Pia Miglietta

## Abstract

The Upside-Down Jellyfish, *Cassiopea*, has become a mainstay of cnidarian photosymbiosis research. Two nominal sister species, *C. xamachana* and the globally introduced *C. andromeda*, supply most of the medusae used in American and European laboratory research within this genus. As founder identity can shape experimental outcomes, here we utilize whole genome resequencing of 21 *Cassiopea* medusae spanning the Florida Keys, Bocas del Toro (Panama), and a European laboratory line, to ask whether mitochondrial species assignment predicts nuclear genome identity. Across multiple population structure analyses using the nuclear genome, Floridian *Cassiopea* carrying *C. xamachana* or *C. andromeda* mitotypes are indistinguishable, and geography is the dominant axis of nuclear genetic structure. A population tree that groups the two Floridian mitotypes as a single interbreeding unit is strongly supported (Patterson’s *D* ≈ 0.0, *Z* = 0.02), whereas a tree that respects mitochondrial species boundaries is rejected (*D* = 0.42, *Z* = 20.8). Strikingly, the European “true” *C. andromeda* line clusters with Panamanian *C. xamachana* rather than with Floridian *C. andromeda*-mitotype animals. From the same sequencing effort, we recover evidence of symbiont variability (*Cladocopium*) in Panama and assemble two near-complete *Tenacibaculum* and *Endozoicomonas* metagenomically-assembled genomes from Floridian host tissue. Together these results indicate that the *C. andromeda/C. xamachana* hybridization zone may extend across ocean basins, and that a “pure” original of either species may be difficult to find. We urge *Cassiopea* researchers to establish new European lines with described genomes.

## Introduction

The Upside-Down Jellyfish, *Cassiopea*, are endemic to tropical and subtropical coastlines worldwide^1^. Over the last decade, *C. andromeda* Forskål, 1775^2^ and *C. xamachana* Bigelow, 1892^3^ have emerged as a cnidarian photosymbiotic model system, used for research on sleep, heat stress, nutrient availability, and more^4–8^. As in other model systems, the choice of starting culture can carry profound implications for experimental outcomes^9,10^. If two nominal laboratory *Cassiopea* species differ in ecology or biology, then identifying lineage boundaries is of the utmost importance.

*Cassiopea* is a genus of striking phenotypic plasticity, yet species determination is restricted to a small number of diagnostic features^11^. Many relationships within this clade have long remained unresolved^12–14^. Phylogenetic surveys of the group recover consistent signals of introduction and crypsis, and the number of undescribed lineages may exceed the number of named species^11,13,14^. This combination of morphological ambiguity, hidden genetic diversity, and extensive human-mediated dispersal complicates efforts to define species boundaries and reconstruct the geographic origins of contemporary populations.

Within this clade, several lines of evidence indicate that *Cassiopea andromeda* has expanded rapidly beyond its presumed native range, likely aided by maritime traffic. Mitochondrial lineages assigned to *C. andromeda* now occur in South America, the Mediterranean, and the Pacific^11,13^. Such introductions can bring previously isolated lineages into secondary contact, creating opportunities for hybridization and introgression. Under these circumstances, mitochondrial markers alone may provide an incomplete or misleading account of evolutionary history. Discordance between mitochondrial identity and nuclear ancestry can arise through recent hybridization, mitochondrial introgression, incomplete lineage sorting, sex-biased dispersal, or repeated introductions from genetically differentiated source populations.

Genome-wide nuclear data are therefore needed to determine whether mitochondrial lineages correspond to reproductively distinct species.

The nominal sister species of *Cassiopea andromeda, C. xamachana*, was described by Bigelow (1892)^3^ on the basis of its distinctive appearance and geographic occurrence, and has since experienced a similarly tangled history of repeated introductions outside of its native range^9,14–16^. Histories of these two species overlap in space and time; in the Florida Keys, *C. xamachana* and a non-native *C. andromeda* lineage co-occur in the same shallow-water assemblages^16^. The Keys therefore represent an important contact zone in which to test whether mitochondrial species assignments correspond to distinct nuclear populations or whether local medusae constitute a shared, potentially admixed gene pool.

The uncertainty around these two species identities has practical consequences. The Florida Keys supply a substantial proportion of the *Cassiopea* stock used in U.S. research and the aquarium trade. If outwardly similar animals collected from the region represent ecologically or physiologically distinct lineages, unrecognized differences in stock identity could reduce the cross-study comparability of laboratory experiments. Alternatively, if mitochondrial lineages are embedded within the same regional nuclear population, mitochondrial species labels may overstate the biological differences among laboratory stocks. Despite the growing use of *Cassiopea* as an experimental system, the population-genomic structure of the Florida contact zone has not previously been characterized.

Here, we use whole-genome resequencing of medusae from Florida, Panama, and laboratory cultures to characterize nuclear divergence, admixture, and geographic structure among lineages assigned to *Cassiopea andromeda* and *C. xamachana*. We ask whether mitochondrial species identity predicts genome-wide nuclear ancestry, whether Floridian mitochondrial lineages form genetically distinct populations, and whether nuclear differentiation is structured more strongly by geography than by mitotype. If these nominal species represent reproductively isolated lineages, we expect mitochondrial identity to correspond closely to nuclear population structure. In contrast, a mismatch between mitochondrial and nuclear variation would support historical or ongoing gene flow, mitochondrial introgression, or taxonomic boundaries that do not reflect genome-wide ancestry. We find that mitotype is not a reliable predictor of nuclear ancestry and that geography structures the nuclear genome far more strongly than mitochondrial species identity. We consider how this discordance between mitochondrial and nuclear DNA affects the interpretation of laboratory studies and what it implies for species boundaries within the global *C. andromeda* complex.

## Methods

### Tissue collection

Oral arm tissue was collected from *Cassiopea* specimens from Florida, Panama, and laboratory-raised specimens. Floridian specimens were collected along the length of the Florida Keys in the summer of 2021, as described previously^16^. Initial species identification was performed by checking 16S and COI haplotypes (see Muffett & Miglietta 2023)^14^, yielding a cohort of nine *C. xamachana*- and five *C. andromeda*-mitotype individuals. An additional cohort of six medusae was collected in 2022 from the mangroves abutting the Smithsonian Tropical Research Institute in Bocas del Toro, Panama (all *C. xamachana* mitotype). One *C. andromeda* medusa, derived from a longstanding Aquarium of Genoa laboratory culture, was received in 2023 and sequenced as an outgroup, for 21 individuals in total (Sup. Tab. S1). Tissue samples were preserved in 100% ethanol (Panama & lab) or Dimethyl sulfoxide EDTA salt-saturated solution (Florida) and stored at -80ºC.

### DNA extraction and sequencing

DNA was extracted using QIAGEN DNeasy kits (Germantown, MD, USA) for all Floridian samples and Zymo Research ZymoBIOMICS Quick-DNA kits (Irvine, CA, USA) for Panamanian samples and the laboratory *C. andromeda* line. All 21 samples were submitted to Azenta Life Sciences for library preparation and sequencing on an Illumina NovaSeq (2×150 bp). Nuclear genomes were sequenced to a mean depth of 8–20x and mitochondrial genomes to 36–245x (Sup. Tab. S2).

### Data preprocessing and variant calling

Raw reads were processed to remove adapters (‘ktrim=r’, ‘k=23’, ‘mink=11’, ‘hdist=1’) and PhiX contamination (‘k=31’, ‘hdist=1’) using BBDuk^17^. To remove extraneous sequence, reads were filtered with BBSplit against masked Symbiodiniaceae reference genomes (*Breviolum minutum, Cladocopium goreaui*, and *Symbiodinium microadriaticum*) and the JGI-provided masked human reference genome (‘k=15’, ambiguous behavior: best match). Cleaned reads were error-corrected with Tadpole (‘mode=correct’, ‘k=50’) and then subsampled to a target of 25 million reads per individual to prevent differences in sequencing depth from biasing downstream comparisons.

Reads were mapped to the chromosome-level *Cassiopea xamachana* reference assembly (GenBank GCA_964235115.1; Sup. Tab. 3). Haplotype and variant calling for the nuclear and mitochondrial genomes was executed with snpArcher^18^, which wraps GATK HaplotypeCaller^19^, using ‘minNmer = 500’, ‘ploidy = 2’, ‘minP = 2’, ‘minD = 4’, ‘het_prior = 0.05’, ‘mappability_min = 1’, and ‘cov_merge = 100’. Before filtering, 20,079,362 SNPs were identified across the 21 samples—roughly 6.6% of the estimated genome size. Variant filtering was performed with VCFtools v0.1.16 (‘--max-alleles 2’, ‘--maxDP 28’, ‘--max-meanDP 28’, ‘-- minDP 10’, ‘--min-meanDP 10’, ‘--minQ 30’, ‘--max-missing 0.9’)^20^. Because of high missingness and low overall depth, one Floridian individual was removed, leaving 20 samples. Of the 20,079,362 sites, 2,470,531 were retained in the unpruned dataset and 1,851,566 after minor-allele-frequency filtration (MAF ≥ 0.08). PLINK v1.9^21^ was used to prune for linkage disequilibrium (‘--indep-pairwise 50 10 0.1’) (Sup. Tab. S4).

### 18S and mitogenome identity

18S rRNA genes were predicted from each *Cassiopea* genome assembly (assembled with SPAdes under standard parameters) with Barrnap v0.9 (kingdom “Euk”) and combined with curated GenBank references (*C. xamachana* AY920771.1; *C. andromeda* HM194818.1 and KY610763–KY610768.1; *C. frondosa* HM194819.1; *C. ornata* HM194785.1; *Cassiopea* sp. AF099675.1) and a *Cephea cephea* outgroup (HM194769.1). In order to confirm predicted 18S sequences, an additional two 18S sequences amplified using the primers and protocol in Abboud et al.^12^ and Sanger sequenced at Texas A&M University Corpus Christi genomics core were included for comparison. An alignment across all sequences was produced with FAMSA2, then trimmed and a tree was created using IQtree with automatic model selection (TN+F+G4) and supported through bootstraps (UltraFast, 1000) and SH-aLRT branch test (1000). Assembly of 18S using Barrnap resulted in a likely artificial subclade not supported by the Sanger sequences.

Mitogenomes were extracted using GetOrganelle v1.7 with the *C. xamachana* mitogenome as a seed (get_organelle_from_reads -F animal_mt -s OZ174019.1 -R 15 -k 21,45,65,85,105 --max-reads 20000000), aligned with FAMSA2 and a maximum likelihood phylogeny was produced using IQtree with automatic model selection (TIM+F+R2) and supported through bootstraps (UltraFast, 1000) and SH-aLRT branch test (1000).

### Population genomics and introgression analysis

To assess population structure and identify putative species boundaries, we performed several complementary clustering and population-genomic analyses. Principal component analysis (PCA) was conducted in PLINK using the filtered set of biallelic SNPs. Individual ancestry proportions were estimated with ADMIXTURE^22^ for *K* = 2–5, with the optimal *K* selected by cross-validation error. Genome-wide genetic differentiation was estimated as Wright’s *F*_ST_ in PLINK across the four sampled groups (Floridian *Cassiopea xamachana* mitotype, Floridian *C. andromeda* mitotype, Panamanian *C. xamachana*, and the European *C. andromeda* line). To test for introgression and to quantify admixture proportions, we computed Patterson’s *D*-statistics (ABBA-BABA tests)^23,24^ with Dsuite trio and lab *C. andromeda* as an outgroup^25^, evaluating both a geography-based population tree and a mitotype-based species tree, and mapped genome-wide admixture signals to branches using the *f*-branch statistic. Individual heterozygosity and inbreeding coefficients (*F*) were estimated in VCFtools.

### Non-*Cassiopea* read identity

Reads that did not align to the *Cassiopea xamachana* reference were classified with Kaiju v1.9^26^ on the KBase platform^27^ against the NCBI BLAST nr database (minimum genus prevalence 0.3%). Genus-level Kaiju output was imported into phyloseq^28^, filtered to remove *Saccharomyces* reads (a likely contaminant of the ethanol used to store the Panamanian samples), and partitioned into Symbiodiniaceae and non-Symbiodiniaceae datasets.

### Metagenome-assembled genome (MAG) recovery and characterization

As several tissue-derived samples carried high abundances of *Tenacibaculum* and *Endozoicomonas* reads, non-*Cassiopea* reads were assembled and co-binned using Metagenome-Atlas^29^. In short, reads were assembled with SPAdes v4.0^30^, then co-binned with VAMB v3.0.2^31^. Assembled bins were assessed for completeness and contamination (CheckM2 v1.0.1^32^) and the two resultant high-quality bins were annotated for coding sequences (DRAM)^33^ and metabolic pathway completeness (KEGG)^34,35^. Secretion systems and ARG profiles were characterizad using PATRIC^36^. N50 was checked with Quast v4.4^37^. Recovered MAGs were compared against host-tissue derived *Endozoicomonas* and *Tenacibaculum* genomes to identify host-association gene repertoires and metabolic capacities using DRAM and core gene trees (AddSpeciestoGenomeTree) within KBase^27^. See Sup. Tab. S5 for the reference genomes used in the comparative analyses.

## Results

### Read mapping and symbiont composition

*Cassiopea* read mapping rates were variable across individuals, with outlier samples having more than 30% of reads map to *Symbiodinium microadriaticum* (sample S117, 66.3%; sample S87, 38.0%; Sup. Tab. S6). Despite this variability, mean mapping rates to the *Cassiopea xamachana* reference were nearly identical for the two mitotypes; 84.3% for *C. andromeda*-mitotype animals (including the laboratory line) and 84.6% for *C. xamachana*-mitotype animals (Sup. Tab. S7).

After adjusting for the proportion of host reads, *S. microadriaticum* was the most-mapped symbiont reference in both the Florida Keys and Panama. The two regions differed in their algal signatures, however: relative to non-host reads, *Cladocopium goreaui* reads appeared roughly ten times more often in Panamanian animals than in Floridian ones, while *Symbiodinium* mapping was about four times lower in Panama (Figure 1a).

**Figure 1.**
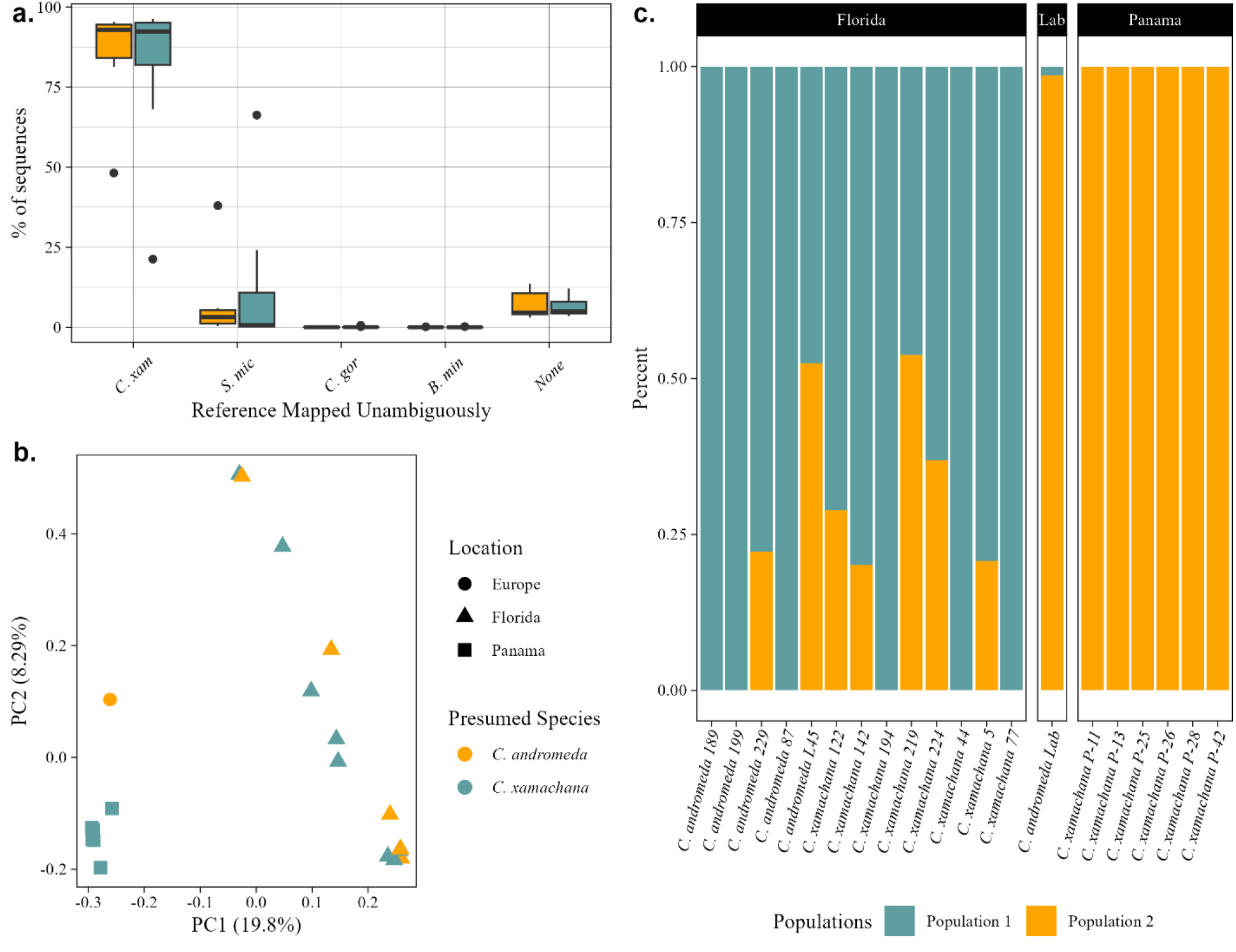
Location rather than mitotype predicts nuclear SNPs. **a**, Raw read mapping rates to the *Cassiopea xamachana* host reference and to three Symbiodiniaceae references, separated by mitochondrial haplotype. Host mapping is near-identical between mitotypes (mean 84.3% vs 84.6%); symbiont mapping is dominated by *S. microadriaticum* and is highly variable among individuals. **b**, Principal component analysis of nuclear SNPs. Points are colored by presumed (mitotype-based) species and shaped by collection location. Nuclear variation is structured by geography, not by mitotype. **c**, ADMIXTURE ancestry proportions at *K* = 2, faceted by collection location. Floridian animals of both mitotypes share a single cluster; Panamanian *C. xamachana* and the European *C. andromeda* line share the other.

### Mito-nuclear discordance and population structure

Putative mitogenomic identity was not indicative of nuclear identity (Figures 1b, SF1 & SF2): Floridian animals carrying the *Cassiopea andromeda* mitotype occupied the same nuclear space as sympatric Floridian *C. xamachana*, and the dominant axis of variation separated Floridian from non-Floridian individuals rather than *C. andromeda* from *C. xamachana* (Sup. Tab. S8). ADMIXTURE largely recovered the same signal. Cross-validation error was minimized at *K* = 2, inferring two clusters corresponding to geography, not to nominal species. One cluster comprised Floridian *Cassiopea* of both mitotypes; the other comprised Panamanian *C. xamachana* together with the European *C. andromeda* line (Figure 1c; Sup. Tab. S9 & S10).

Several Floridian individuals showed mixed ancestry between the two clusters, consistent with an admixed resident population, whereas Panamanian animals were assigned to their cluster essentially without admixture (ancestry ≥ 0.99999). Against expectations, the European laboratory *C. andromeda*, sequenced as a putatively “true” representative of that species, was assigned 98.6% to the Panamanian *C. xamachana* cluster and only 1.4% to the Floridian cluster.

### Population differentiation and tests for gene flow

Genome-wide differentiation among all four groups (*Cassiopea xamachana* Florida, *Cassiopea andromeda* Florida, *Cassiopea andromeda* lab and *Cassiopea xamachana* Panama) was modest (weighted mean *F*_ST_ = 0.11); however comparing Floridian to Panamanian medusae demonstrated clearer partitioning (weighted mean *F*_ST_ = 0.17) and nominal *Cassiopea xamachana* vs *C. andromeda* within Florida demonstrated no barriers (weighted mean *F*_ST_ = - 0.004). ADMIXTURE estimated an *F*_ST_ of 0.25 between its two inferred ancestral populations (log-likelihood -1,306,972 at *K* = 2), placing most of the genome’s structure along the Florida-Panama axis (Sup. Tab. S11).

Formal tests for gene flow found no signal of mitochondrial identity in population definition. When the four groups were arranged according to mitochondrial species identity (Floridian *Cassiopea andromeda* vs. Floridian and Panamanian *C. xamachana)*, Patterson’s *D* was large and highly significant (*D* = 0.42, *Z* = 20.8, *p* ≈ 0). When location-derived topology was tested, grouping all Floridian *Cassiopea* as a single unit relative to Panama and the outgroup, *D* was close to zero (*D* = 0.0003, *Z* = 0.017, *p* = 0.99), suggesting the Floridian mitotypes behave as a single interbreeding population (Sup. Tab. S12 & S13). While these data demonstrate that some barriers between geographically separate *Cassiopea* populations exist, they are not associated with mitochondrial lineage.

### Genetic diversity

Individual inbreeding coefficients were consistent with the population-structure results. Panamanian *Cassiopea xamachana*, compared against a Floridian reference genome, showed uniformly elevated *F* (0.14-0.23). Floridian animals of both mitotypes were more heterozygous on average and more variable (*F* = 0.03-0.32), and one Floridian *C. andromeda*-mitotype individual (L45) carried a slight excess of heterozygosity (*F* = -0.04) consistent with admixed ancestry (Sup. Tab. S14).

### *Cassiopea* includes a specialist *Endozoicomonas* symbiont

The minor taxa recovered from *Cassiopea* tissue do not constitute an unbiased microbiome survey, but they are informative. They support *Cladocopium* as a repeated symbiont of *C. xamachana* in Bocas del Toro, consistent with the elevated *C. goreaui* mapping in Panamanian samples (Figure 2a). In agreement with earlier microbiome work on this system, *Endozoicomonas* was a prominent component of reads from within *Cassiopea* tissue, with *Tenacibaculum* and *Sneathiella* the next most common (Figure 2b)^38,39^. The laboratory-acquired *C. andromeda* harbored only reads assigned to *Rhizophagus*, a fungal genus with no described marine lineages, underscoring the poor transition of native microbial community to long-term culture. Within Floridian tissue samples, *Endozoicomonas* and *Tenacibaculum* were abundant enough to support the assembly of near-complete metagenomically-assembled genomes; in the most extreme case, *Endozoicomonas* reads made up 5% of all DNA recovered from a host tissue section, signaling substantial in-tissue prevalence (Sup. Tab. S15).

**Fig 2.**
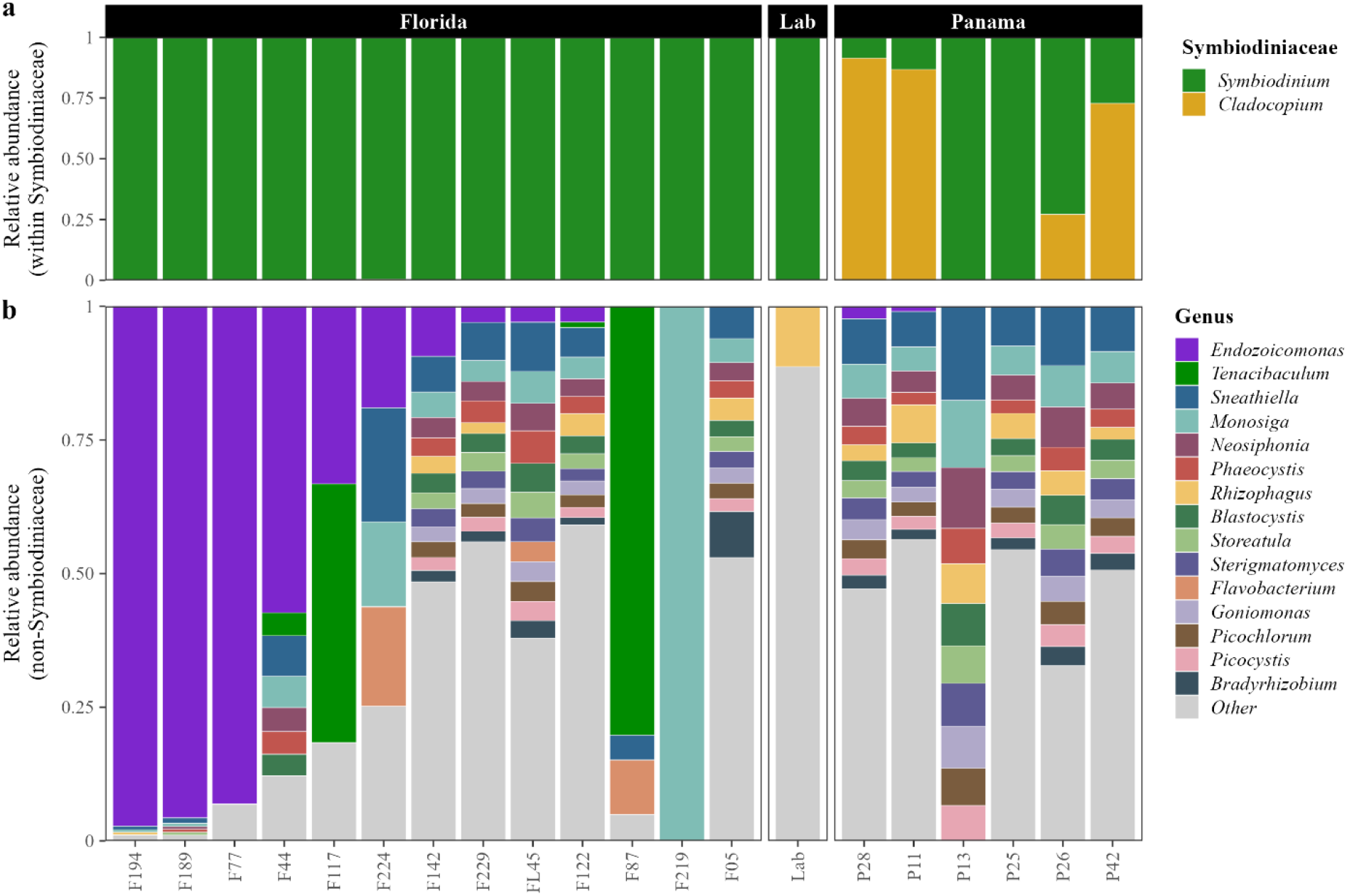
*Endozoicomonas* dominates bacterial reads from Floridian tissues. **a**, Symbiodiniaceae read abundance by genus and **b**, host-tissue derived non-algal reads from Kaiju.

From the non-*Cassiopea* reads we recovered two near-complete MAGs: a *Tenacibaculum* (~2.8 Mb, 2,658 predicted CDS, 99.86% completeness / 0.33% contamination) and an *Endozoicomonas* (~5.2 Mb, 5,262 predicted CDS, 99.99% completeness / 1.36% contamination) (Sup. Tab. S16). The *Endozoicomonas* MAG is metabolically rich, with 78 complete KEGG pathways, including complete galactose degradation, glyoxylate cycling, and methylcitrate cycling (Sup. Tab. S17). Notably, it lacks the DMSP-degradation machinery reported for some coral-associated *Endozoicomonas*^40^, yet it carries a substantial symbiosis- or parasitism-associated gene repertoire, including type IV pili and components of a type III secretion (injectisome) system, features associated with host colonization and host-cell interaction (Sup. Tab. S18). The *Tenacibaculum* MAG encodes a different suite of host-interaction genes, including a complete Type IX secretion / gliding-motility apparatus (T9SS + GldA–N), 13 SusC/SusD polysaccharide-utilization loci indicative of mucus-glycan foraging, and two GH3-family acyl-amido synthetase paralogs whose closest characterized homologs modify indolic and salicylic acids (Sup. Tab. S18, S19). Unlike the coral-associated members of the proposed *Neoendozoicomonas*^41^, this *Endozoicomonas* MAG rests squarely within a growing group of jellyfish-isolated *Endozoicomonas* lineages: it forms a subclade with *Endozoicomonas* from other jellyfish hosts (*Aurelia, Pseudorhiza*) and is sister to a clade containing *E. atrinae* from *Cassiopea ornata*; their function within host tissues remains unidentified (Figure 3).

**Fig 3.**
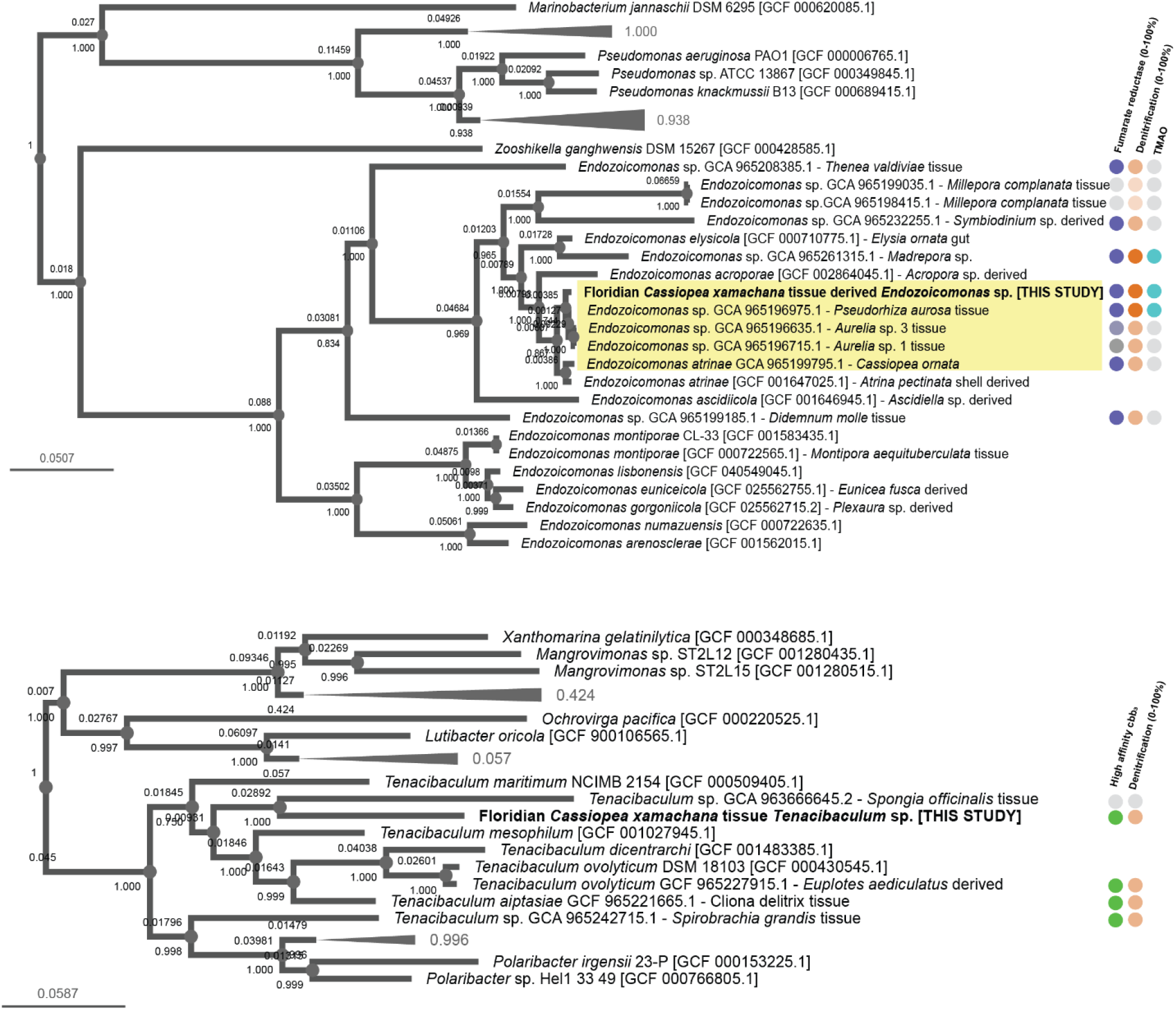
*Endozoicomonas* MAG rests within a scyphozoan-associated clade. Species trees of *Endozoicomonas* (top) and *Tenacibaculum* (bottom) with relative genomic signature of low oxygen acclimation within groups. For *Endozoicomonas*, hypoxia-associated genes include Fumarate pathway completeness (0 (grey) -1 (dark purple)), denitrification steps present (0% (grey)-75% (dark orange)) and TMAO reductase (presence teal, absence grey). For *Tenacibaculum*, this includes Complex IV Cytochrome C cbb_3_-type pathway (presence green, absence grey) and denitrification steps present (0% (grey)-75% (dark orange)).

Interestingly, both MAGs are enriched for microaerobic pathways suited to a low oxygen tissue environment. *Cassiopea* mesoglea have shown a strong diel oxygen oscillation, as low as~28% air saturation before dawn and peaking over 250% air saturation during daylight^8^. The *Tenacibaculum* MAG has a single high-affinity terminal oxidase (cbb_3_-type operon; Sup. Tab. S20, S21), a configuration diagnostic of obligate or near-obligate microaerobes^42,43^. The *Endozoicomonas* MAG includes TMAO reductase (torA-type), fumarate reductase (frdABCD) and denitrification (nitrate to nitrous oxide; Sup. Tab. S20, S22)^44^.

## Discussion

The past two decades have seen a steady contraction in the number of taxa regarded as truly cosmopolitan^45–47^. Among marine invertebrates especially, molecular tools have repeatedly split apparently widespread species into well-defined clades^11,48^. However, the same timeframe has been characterized by massive dissemination of invasive and nonnative coastal invertebrates through shipping and trade^49–51^. Many “distributions” now on record postdate European colonization and the disturbances that came with it^50^. Ecological memory is short, and the true pre-industrial ranges of animals like *Cassiopea* may never have entered the scientific record. Our findings demonstrate that, in the Florida Keys, genetic divergence of 7% between the *C. andromeda* and *C. xamachana* mitochondrial genomes is not accompanied by corresponding nuclear genomic differentiation^11^. The co-occuring animals of the two mitotypes represent a single interbreeding population. Within our data, *C. andromeda* and *C. xamachana* are not differentiable under a biological species concept^52^. If this holds with further investigation, *Cassiopea andromeda*, being the senior synonym, takes priority. The nuclear genome of the European “true” *C. andromeda* line clusters with Panamanian *C. xamachana*, a result that requires additional genome-wide validation in a more comprehensive global dataset. Based on these data alone, the hybridization or connectivity zone of the *C. andromeda* species complex may extend across ocean basins, and truly “pure” lineages of either nominal species, if they persist, may be the exception rather than the rule. We cannot confirm the full geographic extent of this connectivity from three locales, testing of the extent of this requires broad global sampling.

These results carry a concrete cautionary message for the *Cassiopea* research community. Laboratory cultures sourced from Florida are routinely treated as *C. xamachana*, and animals are moved between laboratories and mitotypes on the assumption that mitochondrial barcodes equate to genomic identity. Our results indicate that COI or 16S haplotype is a poor predictor of nuclear ancestry in Floridian individuals. Consequently, founder colonies assigned to different species may be indistinguishable at the nuclear level, whereas colonies assigned to the same species but originating from different regions may exhibit substantial nuclear differentiation. Based on these data, we call on the establishment of additional complementary European lines for lab studies, with genome sequencing comparable to the US T1-A line^53^. More broadly, if complete nuclear mixing is compatible with this degree of mitochondrial divergence in *Cassiopea*, then population-genomic surveys of other understudied cnidarians are warranted, as loss of genetic diversity in admixed or introduced populations can carry consequences for fitness and for the ecosystem services these animals provide.

Finally, the excess reads captured by whole-genome sequencing yielded important ecological information. We found support for a regional relationship with Cladocopium in Bocas del Toro medusae. As symbionts have profound effects on host physiology, this diversity in native symbionts provides new potential study sites^54,55^. We also recovered host-specialized hypoxia-adapted bacterial associates; the *Endozoicomonas* MAG, abundant enough in one host to comprise 5% of recovered DNA, appears to lack DMSP machinery but retains a host-interaction toolkit (type IV pili, a type III injectisome); whether it is mutualist, commensal, or pathobiont in *Cassiopea* remains unresolved. The clustering of jellyfish-associated *Endozoicomonas* MAGs provides a new lens for *Endozoicomonas* coevolution beyond coral, suggesting that *Endozoicomonas* may include a scyphozoan-associated clade sister to *E. atrinae*. As *Endozoicomonas acroporae* tissue aggregates have newly discovered benefits for host protein folding stability under heat stress, *Endozoicomonas* may provide environmental hardening within *Cassiopea* as well^56^. Together with the *Tenacibaculum* genome, these provide new additional resources for those interested in bacterial aggregates within cnidarian host tissues.

## Supporting information

Sfigs 1&2

## Data and code availability

All sequences associated with this work are available via accession PRJEB111638.

## References

1. Ohdera, A. H. et al. Upside-down but headed in the right direction: Review of the highly versatile Cassiopea xamachana system. Front. Ecol. Evol. 6, 1–15 (2018).

2. P., F. Descriptiones Animalium, Avium, Amphibiorum, Piscium, Insectorum, Vermium; quae in Itinere Orientali Observavit Petrus Forskål. (1775).

3. Bigelow, R. P. On a new species of Cassiopea from Jamaica. Zool. Anz. 15, 212–214. (1892).

4. Hofmann, D. K., Neumann, R. & Henne, K. Strobilation, budding and initiation of scyphistoma morphogenesis in the rhizostome Cassiopea andromeda (Cnidaria: Scyphozoa). Mar. Biol. 47, 161–176 (1978).

5. McGill, C. J. & Pomory, C. M. Effects of bleaching and nutrient supplementation on wet weight in the jellyfish Cassiopea xamachana (Bigelow) (Cnidaria: Scyphozoa). Mar. Freshw. Behav. Physiol. 41, 179–189 (2008).

6. Nath, R. D. et al. The Jellyfish Cassiopea Exhibits a Sleep-like State. Curr. Biol. 27, 2984–2990.e3 (2017).

7. Banha, T. N. S., Mies, M., Güth, A. Z., Pomory, C. M. & Sumida, P. Y. G. Juvenile Cassiopea andromeda medusae are resistant to multiple thermal stress events. Mar. Biol. 167, 1–13 (2020).

8. Arossa, S. et al. The Internal Microenvironment of the Symbiotic Jellyfish Cassiopea sp. From the Red Sea. Front. Mar. Sci. 8, 1–11 (2021).

9. Bellis, E. S. & Denver, D. R. Natural variation in responses to acute heat and cold stress in a sea anemone model system for coral bleaching. Biol. Bull. 233, 168–181 (2017).

10. Suzuki, T. A. et al. Selection and transmission of the gut microbiome alone can shift mammalian behavior. Nat. Commun. 16, (2025).

11. Mora, E. G., Collins, A. G., Boco, S. R., Geson, S. M. & Morandini, A. C. Revealing hidden diversity among upside-down jellyfishes (Cnidaria: Scyphozoa: Rhizostomeae: Cassiopea): distinct evidence allows the change of status of a neglected variety and the description of a new species. Invertebr. Syst. 36, 63–89 (2022).

12. Schembri, P. J., Deidun, A. & Vella, P. J. First record of Cassiopea andromeda (Scyphozoa: Rhizostomeae: Cassiopeidae) from the central Mediterranean Sea. Mar. Biodivers. Rec. 3, 1–2 (2010).

13. Holland, B. S., Dawson, M. N., Crow, G. L. & Hofmann, D. K. Global phylogeography of Cassiopea (Scyphozoa: Rhizostomeae): Molecular evidence for cryptic species and multiple invasions of the Hawaiian Islands. Mar. Biol. 145, 1119–1128 (2004).

14. Abboud, S. S., Daglio, L. G. & Dawson, M. N. A global estimate of genetic and geographic differentiation in macromedusae-implications for identifying the causes of jellyfish blooms. Mar. Ecol. Prog. Ser. 591, 199–216 (2018).

15. Stampar, S. N. et al. The puzzling occurrence of the upside-down jellyfish cassiopea (Cnidaria: Scyphozoa) along the Brazilian coast: A result of several invasion events? Zoologia 37, 1–10 (2020).

16. Muffett, K. & Miglietta, M. P. Demystifying Cassiopea species identity in the Florida Keys: Cassiopea xamachana and Cassiopea andromeda coexist in shallow waters. PLoS One 18, 1–15 (2023).

17. Bushnell, B. BBMap: A Fast, Accurate, Splice-Aware Aligner. in USDOE vol. 13 1–2 (2012).

18. Mirchandani, C. D. et al. A Fast, Reproducible, High-Throughput Variant Calling Workflow for Population Genomics. Mol. Biol. Evol. 41, 1–15 (2024).

19. McKenna, A. et al. The genome analysis toolkit: A MapReduce framework for analyzing next-generation DNA sequencing data. Genome Res. 20, 1297–1303 (2010).

20. Danecek, P. et al. The variant call format and VCFtools. Bioinformatics 27, 2156–2158 (2011).

21. Purcell, S. et al. PLINK: A tool set for whole-genome association and population-based linkage analyses. Am. J. Hum. Genet. 81, 559–575 (2007).

22. Alexander, D. H., Novembre, J. & Lange, K. Fast model-based estimation of ancestry in unrelated individuals. Genome Res. 19, 1655–1664 (2009).

23. Patterson, N. et al. Ancient admixture in human history. Genetics 192, 1065–1093 (2012).

24. Durand, E. Y., Patterson, N., Reich, D. & Slatkin, M. Testing for ancient admixture between closely related populations. Mol. Biol. Evol. 28, 2239–2252 (2011).

25. Malinsky, M., Matschiner, M. & Svardal, H. Dsuite - Fast D-statistics and related admixture evidence from VCF files. Mol. Ecol. Resour. 21, 584–595 (2021).

26. Menzel, P., Ng, K. L. & Krogh, A. Fast and sensitive taxonomic classification for metagenomics with Kaiju. Nat. Commun. 7, (2016).

27. Arkin, A. P. et al. KBase: The United States department of energy systems biology knowledgebase. Nat. Biotechnol. 36, 566–569 (2018).

28. McMurdie, P. J. & Holmes, S. Phyloseq: An R Package for Reproducible Interactive Analysis and Graphics of Microbiome Census Data. PLoS One 8, (2013).

29. Kieser, S., Brown, J., Zdobnov, E. M., Trajkovski, M. & McCue, L. A. ATLAS: A Snakemake workflow for assembly, annotation, and genomic binning of metagenome sequence data. BMC Bioinformatics 21, 1–8 (2020).

30. Bankevich, A. et al. SPAdes: A new genome assembly algorithm and its applications to single-cell sequencing. J. Comput. Biol. 19, 455–477 (2012).

31. Nissen, J. N. et al. Improved metagenome binning and assembly using deep variational autoencoders. Nat. Biotechnol. 39, 555–560 (2021).

32. Chklovski, A., Parks, D. H., Woodcroft, B. J. & Tyson, G. W. CheckM2: a rapid, scalable and accurate tool for assessing microbial genome quality using machine learning. Nat. Methods 20, 1203–1212 (2023).

33. Shaffer, M. et al. DRAM for distilling microbial metabolism to automate the curation of microbiome function. Nucleic Acids Res. 48, 8883–8900 (2020).

34. Kanehisa, M., Furumichi, M., Sato, Y., Kawashima, M. & Ishiguro-Watanabe, M. KEGG for taxonomy-based analysis of pathways and genomes. Nucleic Acids Res. 51, D587–D592 (2023).

35. Kanehisa, M., Sato, Y. & Morishima, K. BlastKOALA and GhostKOALA: KEGG Tools for Functional Characterization of Genome and Metagenome Sequences. J. Mol. Biol. 428, 726–731 (2016).

36. Wattam, A. R. et al. Improvements to PATRIC, the all-bacterial bioinformatics database and analysis resource center. Nucleic Acids Res. 45, D535–D542 (2017).

37. Gurevich, A., Saveliev, V., Vyahhi, N. & Tesler, G. QUAST: Quality assessment tool for genome assemblies. Bioinformatics 29, 1072–1075 (2013).

38. Carabantes, N., Cerqueda-García, D., García-Maldonado, J. Q. & Thomé, P. E. Changes in the Bacterial Community Associated With Experimental Symbiont Loss in the Mucus Layer of Cassiopea xamachana Jellyfish. Front. Mar. Sci. 9, 1–13 (2022).

39. Muffett, K. M., Labonté, J. M. & Miglietta, M. P. Florida Keys Cassiopea host benthos-like external microbiomes and a gut dominated by Vibrio, Endozoicomonas and Mycoplasma. PLoS One 20, 1–22 (2025).

40. Kuek, F. W. I. et al. DMSP Production by Coral-Associated Bacteria. Front. Mar. Sci. 9, 1–12 (2022).

41. Garritano, A. N. et al. Resolving the evolutionary duality of marine symbionts : redefining the genus Endozoicomonas and proposing Neoendozoicomonas gen . nov. 6, 1–12 (2026).

42. Durand, A. et al. Biogenesis of the bacterial cbb3 cytochrome c oxidase: Active subcomplexes support a sequential assembly model. J. Biol. Chem. 293, 808–818 (2018).

43. Appleby, C. A., Preisig, O., Zufferey, R., Tho, L. & Hennecke, H. A High-Affinity cbb 3 - Type Cytochrome Oxidase Terminates the Symbiosis-Specific Respiratory Chain of Bradyrhizobium japonicum. 178, 1532–1538 (1996).

44. Jameson, E. et al. Metagenomic data-mining reveals contrasting microbial populations responsible for trimethylamine formation in human gut and marine ecosystems. Microb. genomics 2, e000080 (2016).

45. Miglietta, M. P., Piraino, S., Kubota, S. & Schuchert, P. Species in the genus Turritopsis (Cnidaria, Hydrozoa): A molecular evaluation. J. Zool. Syst. Evol. Res. 45, 11–19 (2007).

46. Cerca, J., Meyer, C., Purschke, G. & Struck, T. H. Delimitation of cryptic species drastically reduces the geographical ranges of marine interstitial ghost-worms (Stygocapitella; Annelida, Sedentaria). Mol. Phylogenet. Evol. 143, 106663 (2020).

47. Dawson, M. N. & Jacobs, D. K. Molecular evidence for cryptic species of Aurelia aurita (Cnidaria, Scyphozoa). Biol. Bull. 200, 92–96 (2001).

48. Dawson, M. N. & Martin, L. E. Geographic variation and ecological adaptation in Aurelia (Scyphozoa, Semaeostomeae): Some implications from molecular phylogenetics. Hydrobiologia 451, 259–273 (2001).

49. Graham, W. M. & Bayha, K. M. Biological Invasions by Marine Jellyfish. Biol. Invasions 193, 239–255 (2007).

50. Morandini, A. C., Stampar, S. N., Maronna, M. M. & Da Silveira, F. L. All non-indigenous species were introduced recently? the case study of Cassiopea (Cnidaria: Scyphozoa) in Brazilian waters. J. Mar. Biol. Assoc. United Kingdom 97, 321–328 (2017).

51. Maggio, T. et al. Molecular identity of the non-indigenous Cassiopea sp. from Palermo Harbour (central Mediterranean Sea). J. Mar. Biol. Assoc. United Kingdom 99, 1765–1773 (2019).

52. Wheeler, Q. D. & Meier, R. Species concepts and phylogenetic theory: a debate. (Columbia University Press, 2000).

53. Ohdera, A. et al. Box, stalked, and upside-down? Draft genomes from diverse jellyfish (cnidaria, acraspeda) lineages: Alatina alata (cubozoa), calvadosia cruxmelitensis (staurozoa), and cassiopea xamachana (scyphozoa). Gigascience 8, 1–15 (2019).

54. Herrera, M. et al. Unfamiliar partnerships limit cnidarian holobiont acclimation to warming. Glob. Chang. Biol. 26, 5539–5553 (2020).

55. Rodriguez-Casariego, J. A., Cunning, R., Baker, A. C. & Eirin-Lopez, J. M. Symbiont shuffling induces differential DNA methylation responses to thermal stress in the coral Montastraea cavernosa. Mol. Ecol. 31, 588–602 (2022).

56. Lu, C. et al. Endozoicomonas acroporae enhances coral thermal resilience through host – microbe coordination. 20, 1–14 (2026).

