## Supplementary material for "Can an original be found? Mitochondrial species identity does not predict nuclear genome similarity in the photosymbiotic jellyfish *Cassiopea andromeda* and *C. xamachana*": Sfigs 1&2

**Figure SF1. 18S tree of *Cassiopea***

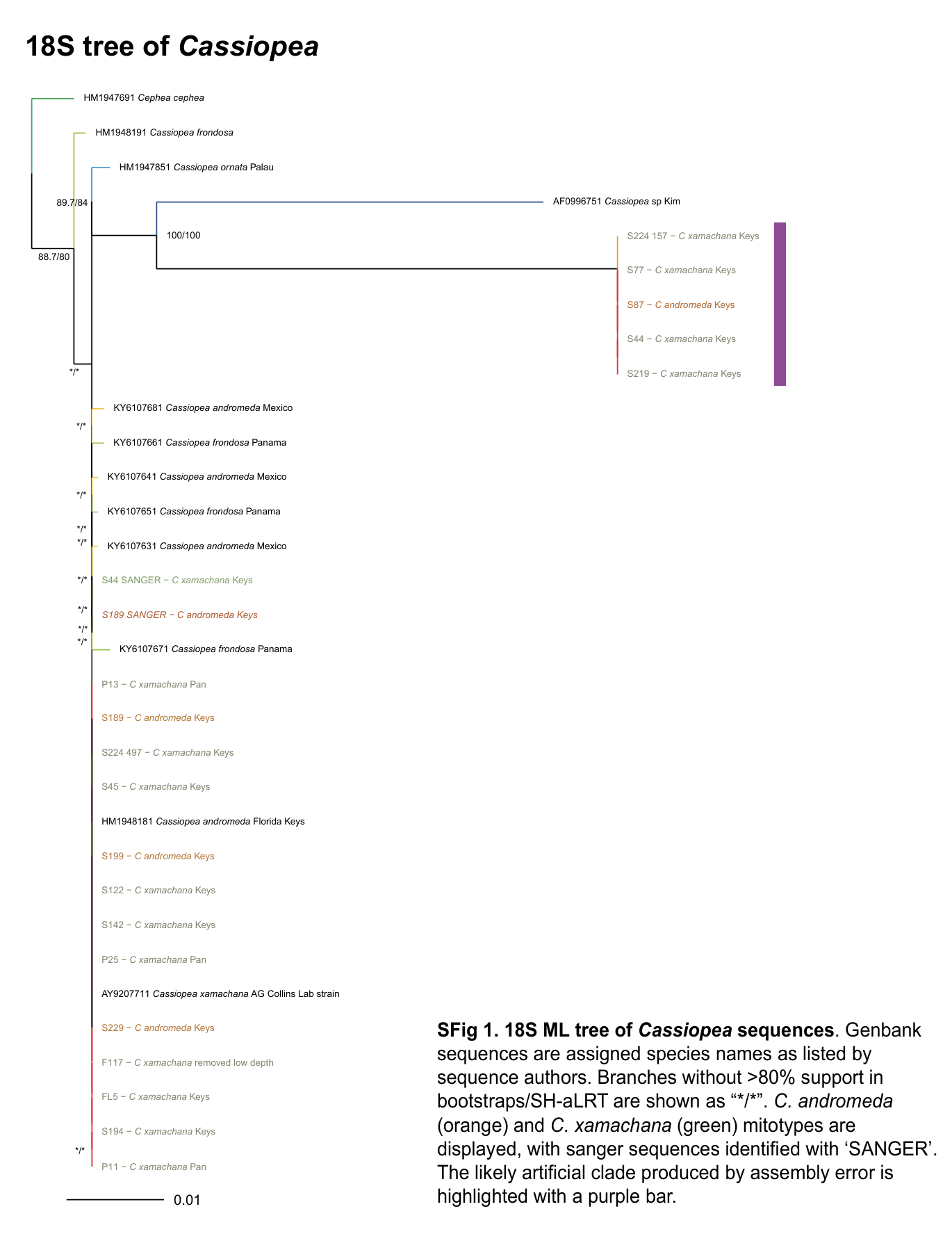

**Figure SF2. ML tree of *Cassiopea* mitogenomes**

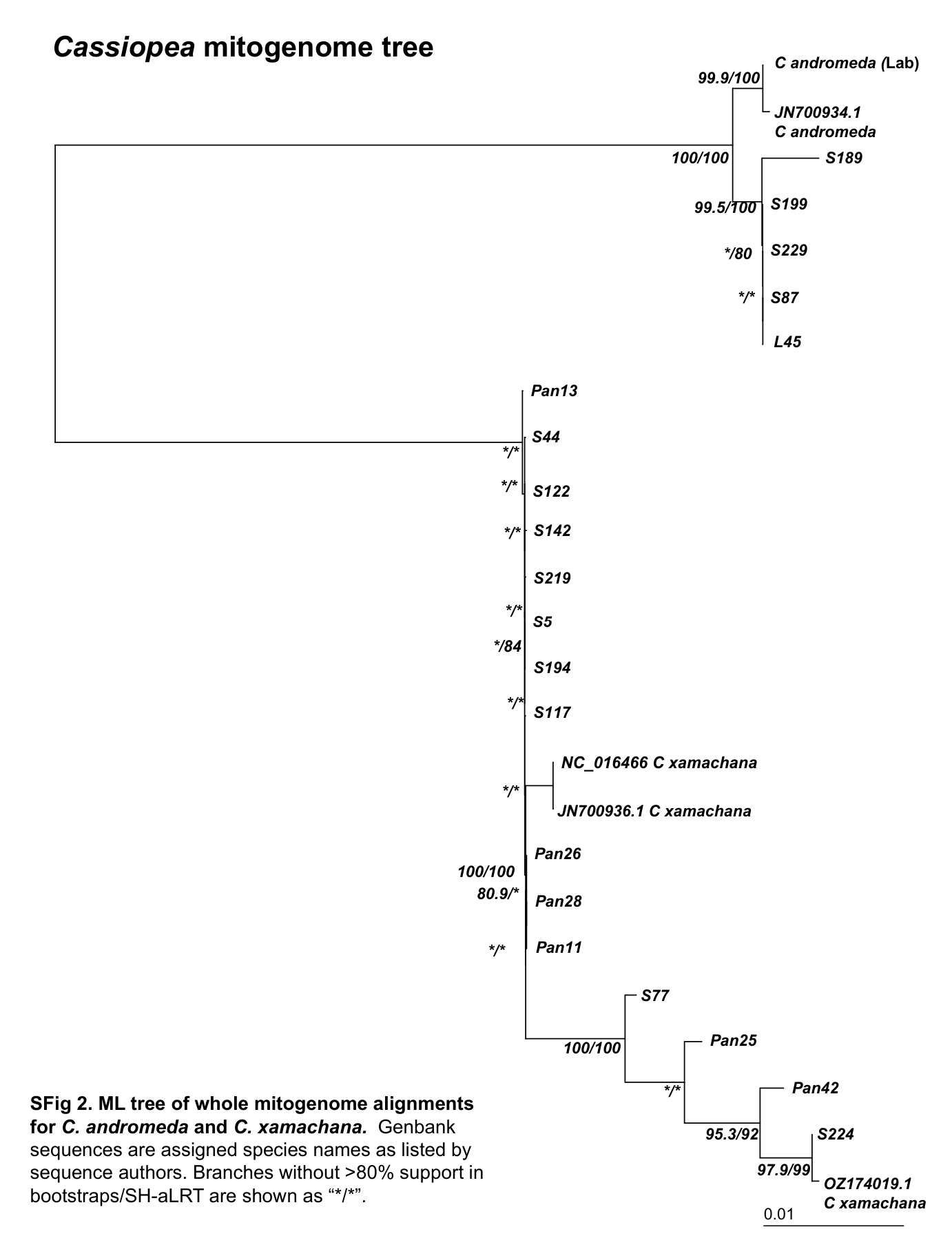
